# Single-Cell Transcriptomics Identifies Dysregulated Metabolic Programs of Aging Alveolar Progenitor Cells in Lung Fibrosis

**DOI:** 10.1101/2020.07.30.227892

**Authors:** Jiurong Liang, Guanling Huang, Xue Liu, Forough Taghavifar, Ningshan Liu, Changfu Yao, Nan Deng, Yizhou Wang, Ankita Burman, Ting Xie, Simon Rowan, Peter Chen, Cory Hogaboam, Barry Stripp, S. Samuel Weigt, John Belperio, William C. Parks, Paul W. Noble, Dianhua Jiang

## Abstract

Aging is a critical risk factor in progressive lung fibrotic diseases such as idiopathic pulmonary fibrosis (IPF). Loss of integrity of type 2 alveolar epithelial cells (AEC2s) is the main causal event in the pathogenesis of IPF. To systematically examine the genomic program changes of AEC2s with aging and lung injury, we performed unbiased single cell RNA-seq analyses of lung epithelial cells from either uninjured or bleomycin-injured young and old mice. Major lung epithelial cell types were readily identified with canonical cell markers in our dataset. Heterogenecity of AEC2s was apparent, and AEC2s were then classified into three subsets according to their gene signatures. Genes related to lipid metabolism and glycolysis were significantly altered within these three clusters of AEC2s, and also affected by aging and lung injury. Importantly, IPF AEC2s showed similar genomic programming and metabolic changes as that of AEC2s from bleomycin injured old mouse lungs relative to controls. Furthermore, perturbation of both lipid metabolism and glycolysis significantly changed progenitor renewal capacity in 3-Demensional organoid culture of AEC2s. Taken togather, this work identified metabolic defects of AEC2s in aging and during lung injury. Strategies to rectify these altered programs would promote AEC2 renewal which in turn improves lung repair.

**One sentence summary:** Metabolic defects of alveolar progenitors in aging and during lung injury impair their renewal.

## INTRODUCTION

Despite extensive efforts, the mechanisms controlling progressive tissue fibrosis remain poorly understood. Growing evidence suggests that idiopathic pulmonary fibrosis (IPF), a fatal form of interstitial lung disease, is a result of repeated epithelial cell injury and inadequate alveolar epithelial repair that leads to excessive fibroblast activity and lung fibrosis (Noble et al., 2012). Type 2 alveolar epithelial cells (AEC2s) function as progenitor cells that maintain epithelium homeostasis and repair damaged epithelium after injury (Barkauskas et al., 2013; Desai et al., 2014; Hogan et al., 2014; Jiang et al., 2020). We and other showed that AEC2s were reduced in the lungs of patients with IPF, and the progenitor cell function of IPF AEC2s was significantly impaired (Liang et al., 2016). In consistent with our observation, a recent study using 3D organoid culture suggests that distal lung epithelial progenitor cell function declines with age (Watson et al., 2020).

The incidence, prevalence, and mortality of IPF all increase with age (Raghu et al., 2016; Rojas et al., 2015). Aging has also been linked to lung fibrosis in animal models (Bueno et al., 2018). Phenotypes of cellular aging including epithelial cell apoptosis (Korfei et al., 2008; Lawson et al., 2011), endoplasmic reticulum stress stress (Burman et al., 2018), autophagy (Sosulski et al., 2015), senescence (Minagawa et al., 2011), telomere shortening (Alder et al., 2008; Naikawadi et al., 2016), mitochondria dysfunction (Bueno et al., 2015; Ryu et al., 2017), oxidative stress (Anathy et al., 2018), and metabolic dysfunction (Liu and Summer, 2019) are observed in IPF lungs. Although the concept that IPF as a disease of aging is well accepted in the field (Martinez et al., 2017; Selman and Pardo, 2014; Thannickal, 2013), there are limited studies focusing on the age-related mechanisms by which aging contributes to the disease development and progression of IPF (Rojas et al., 2015).

The lung is a metabolically active organ. Metabolic dysregulation has been reported in lung epithelial cells from patients with IPF. Proteomics profiling identified the alteration in several metabolic pathways including adenosine triphosphate degradation pathway and glycolysis pathway in IPF lungs (Kang et al., 2016). This would be in line with the observation that IPF exhibit marked accumulation of dysmorphic and dysfunctional mitochondria with reduced expression of PTEN-induced putative kinase 1 (PINK1) (Bueno et al., 2015). These events likely cause apoptosis of AEC2s in patients with IPF (Korfei et al., 2008).

The role of lipid metabolism has been suggeted in IPF and aging (Johnson and Stolzing, 2019; Liu and Summer, 2019). A recent study using single cell transcriptomics and deep tissue proteomics identified dysregulated lipid metabolism in AEC2s during aging (Angelidis et al., 2019). Inhibiting lipid synthesis in AEC2 cells exacerbates bleomycin-induced lung fibrosis in mice (Chung et al., 2019). These studies highlight the crucial role of lipid metobolic dysregulation in the lung during aging and fibrosis. However, the mechanism that regulates cell metabolic reprogramming during aging and fibrosis development is poorly understood.

Much work has been done in fibroblasts showing glucose metabolic dysregulation contributes to progression of lung fibrosis. Lactic acid concentrations are elevated in IPF lung tissue (Kottmann et al., 2012), and lactic acid induces myofibroblast differentiation (Kottmann et al., 2012; Xie et al., 2015), suggesting a glycolysis reprogramming in IPF fibroblasts. Glucose transporter 1–dependent glycolysis is elevated in fibroblasts of aged mouse lungs and contributes to aging related lung fibrosis (Cho et al., 2017), suggesting the profibrosis metabolic alteration is associated with aging. Currently, there is lack of study with glycolysis reprogramming in AEC2s during aging and in lung fibrosis.

In the current study, we took an unbiased approach – single-cell RNA-sequencing (scRNA-seq) – to systematically investigate the genetic signatures and programs of lung epithelial cells in young and old mouse lungs in the static stage and after experimental lung injury, and comparatively examined these programs in IPF lung. We identified three subsets of AEC2s and found the genomic programming changes and metabolic alterations of AEC2 subsets were correlated to lung injury and influenced by aging as well. Most importantly, our data showed that AEC2s of IPF lung have the similar gene signature and metabolic alteration as AEC2s from bleomycin-injured old mouse lungs. Furthermore, we demonstrated that lipid metabolism and glycolysis metabolism regulate alveolar progenitor renewal with 3D organoid culture of primary mouse and human AEC2s.

## RESULTS

### Epithelial cell transcriptome profiles in young and old mouse lungs

To investigate the genetic signatures and programs of lung epithelial cells under the influence of aging and lung injury, we employed an unbiased approach to build an atlas of lung epithelial cells. ScRNA-seq was performed on flow sorted Lin^−^EpCAM^+^ cells from lungs of uninjured as well as bleomycin-injured (day 4 and day 14) young and old mice with 10X Genomics (Figure 1a). The data was aligned with Cell Ranger, and analyzed with Seurat package (Butler et al., 2018). A total of 71,930 cells were further analyzed. Data show that cells from 20 individual mice overlaped very well (Figure 1b).

**Figure 1.**
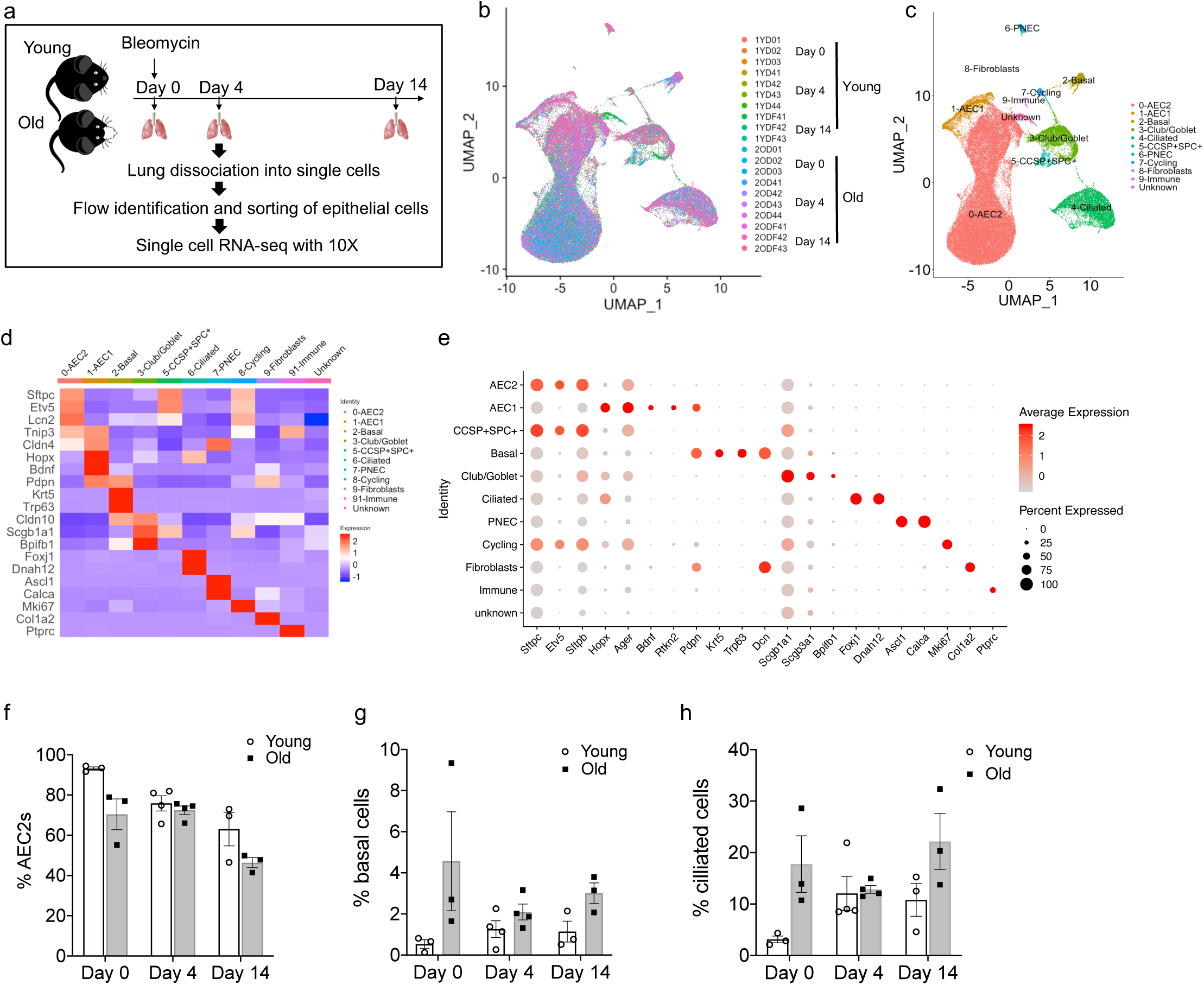
Transcriptome profiles of lung epithelial cells in young and old mice. (a) Schematic of scRNA-seq analysis of flow sorted Lin^−^EpCAM^+^ cells from lungs of uninjured (n = 3), 4 days (n = 4) or 14 dpi (n = 3) young and old mice. (b) UMAP visualization of the cells from all 20 samples. (c) UMAP visualization of epithelial cell clusters. (d) Heatmap of epithelial cell clusters. (e) Dot plots of conventional marker genes of epithelial cell clusters. (f-h). Percentage of AEC2s (g), basal cells (h), and ciliated cells (i) in total epithelial cells (day 0, n = 3; day 4, n = 4; day 14, n = 3).

The major lung epithelial cell types, AEC2, AEC1, basal cell, club cell, ciliated cell, pulmonary neuroendocrine cell (PNEC), and proliferative (cycling) cells were readily identified with canonical cell markers (Figure 1c-e). Since there were only a few cells expressing *Foxi1* scattered in the AEC2 clusters and they did not form a distinct cluster, ionocyte was not identified in the analysis. In addition to the canonical cell markers, novel cell markers for mouse lung epithelial cell types were also identified, but were not reported in this paper since the paper focuses on the heterogeneity of AEC2s.

AEC2 cluster is the largest cluster among the lung epithelial cell populations (Figure 1c,f). Over 90% of total epithelial cells were AEC2s in the uninjured young mouse lungs (Figure 1f). The percentage of AEC2s was reduced in the bleomycin injured lungs. When compared the cells from the lungs of old mouse to the cells from young mice, we observed a trend of decrease of percentage of AEC2s (Figure 1f) along with a trend of increase in basal cells (Figure 1g) and cilliated cells (Figure 1h) within the total epithelial population of old mouse lung relative to that of the young mouse lung at both intact stage and 14 days post injury (dpi). These data are in line with an increase in basal cells in IPF (Adams et al., 2020; Carraro et al., 2020). This result is consistent with a recent report that there was an incrase in cilliated cell population in old mouse lungs (Angelidis et al., 2019).

### Heterogeneity of AEC2s

We hypothesized that the heterogeneity of AEC2s may be influenced by aging and lung injury. Next, we analyzed the gene expressions of a total of 49,571 AEC2s from both homeostatic and bleomycin injuried young and old mouse lungs and identified three AEC2 subsets, AEC2-1, AEC2-2, and AEC2-3 according to their gene expression signatures (Figure 2a, 2b). AEC2-2 showed the lowest correlation silhouette value suggesting its intermediate status between AEC2-1 and AEC2-3 (Figure 2c). Pseudotime analysis identified AEC2-1 with lowest entropy and AEC2-3 with high entropy, suggestive a trasition from AEC2-1 to AEC2-2 and further to AEC2-3. Correlation spanning tree analysis suggested the sequential dynamics (Figure 2d).

**Figure 2.**
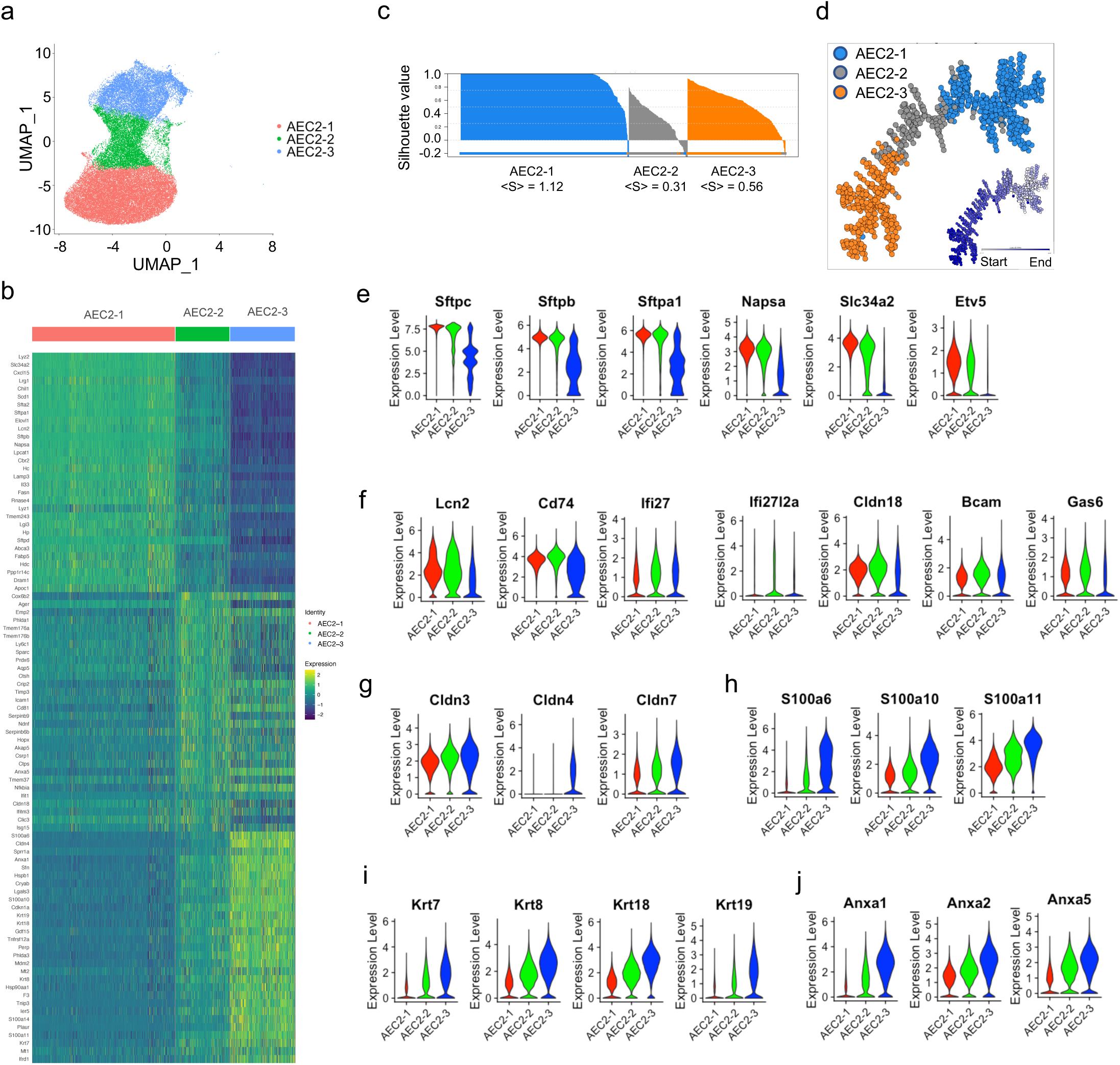
Definition of AEC2 subsets. (a) UMAP of 49,571 AEC2s showing three subsets of AEC2s (red, AEC2-1; green, AEC2-2; and blue, AEC2-3). (b) Heat map representing characteristics of three subsets of AEC2s. Each column represents the average expression value for one cell, grouped by cell cluster. Gene expression values are normalized in rows. (c) Correlation silhouette of AEC2 subsets (d) Correlation spanning tree of AEC2 subsets (e) Violin plots of expression of AEC2-1 markers in all AEC2s (red, AEC2-1; green, AEC2-2; and blue, AEC2-3). (f) Violin plots of expression of AEC2-2 markers in all AEC2s. (g) Violin plots of expression of the claudin family genes in three subsets of AEC2s. (h) Violin plots of expression of the S100 protein family genes in three subsets of AEC2s. (i) Violin plots of expression of the keratin family genes in three subsets of AEC2s. (j) Violin plots of expression of the annexin family genes in three subsets of AEC2s.

Subset AEC2-1 cells express typical signature genes of AEC2s including *Sftpc, Sftpb, Sftpa1, Napsa, Slc34a2*, and *Etv5*. All these AEC2 marker genes were down regulated in cluster 2 and cluster 3 AEC2s (Figure 2e).

Increased interferon signaling was observed in subset AEC2-2 cells. For example, interferon induced genes, *Ifi27* and *Ifi2712a* were upregulated subset AEC2-2 cells (Figure 2f). Other genes up-regulated in subset AEC2-2 cells include *Lcn2, Cd74, Cldn18, Bcam*, and *Gas6* (Figure 2f). The functions of these genes are related to inflammation, epithelial tight junction, MHC class II protein, cell adhension, and growth arrest.

We observed several gene families were up-regulated in subset AEC2-2 cells and their expression were further elevated in subset AEC2-3 cells. These gene families include claudin family (Figure 2g), gene family of heat shock proteins (Figure 2h), keratin family (Figure 2i), and annxin family genes (Figure 2j).

*Cldn4* was only expressed in cluster 3 cells, while *Cldn3* and *Cldn7* expressed on all three cluster of AEC2s with highest expression in cluster 3 (Figure 2g). S100 protein family genes including *S100a6, S100a10*, and *S100a11* were all upregulated in cluster 2 and further elevated in cluster 3 AEC2s (Figure 2h). Several keratin genes (*Krt7, Krt8, Krt18*, and *Krt19*) were upregulated in cluster 2 and further elevated in cluster 3 AEC2s (Figure 2i). The annexin family genes upregulated in cluster 2 and further elevated cluster 3 AEC2s. (Figure 2h).

We further performed pathway analysis of the three subsets of AEC2s. Compared to subset AEC2-1, both subsets AEC2-2 and AEC2-3 showed upregulated oxidative phosphorylation and the p53 signaling. Down regulated EIF2 signaling was obersed in both cluster 2 and cluster 3 cells (Supplementary Table 1, 2). The sirtuin signaling was down-regulated in cluster 2 AEC2s, however it is up-regulated in cluster 3 AEC2s relative to cluster 1 (Supplementary Table 1, 2).

The gene signature and signaling pathways of the three AEC2 subsets indicated that AEC2s in subset AEC2-1 were mainly intact AEC2s, while AEC2s in subset AEC2-2 and subset AEC2-3 were injured AEC2s.

### AEC2 subsets were influenced by both aging and lung injury

Next, we investigated how aging and lung injury affect the subsets of AEC2s by comparing the AEC2s from the lungs of both young and old mice with and without bleomycin injury. Mice at Day 0 were uninjured with intact lung. Day 4 after bleomycin was the timepoint with mixmum AEC2 injury, while 14 dpi with substantial AEC2 recovery (Liang et al., 2016). Over eighty percent of total AEC2s of uninjured young mouse lungs were AEC2-1 cells and the percentage of AEC2-1 cells was slightly lower in the uninjured old mouse lungs (Figure 3a,b). At 4 dpi, AEC2s shifted from subset AEC2-1 to AEC2-2 and AEC2-3 in both young and old mouse lungs. It is interesting that at 14 dpi, the AEC2 recovery stage, the intact AEC2s in AEC2-1 were partially recoved in young mouse lungs, but the old mouse lungs were continuously losing cells in AEC2-1 subset. The majority of AEC2s in old mouse lungs at 14 dpi were subset AEC2-3 cells (Figure 3a,b). These data suggest that the aging affects AEC2 subsets in the lungs and there were reduced subset of intact AEC2s and increased proportion of damaged AEC2s in old mouse lungs especially after injury. The alteration of AEC2 subsets in old mouse lungs after injury relative to the AEC2s from young mouse lungs might be resulted from increased apoptosis and decreased renewal capacity of AEC2s in the aged mouse lungs.

**Figure 3.**
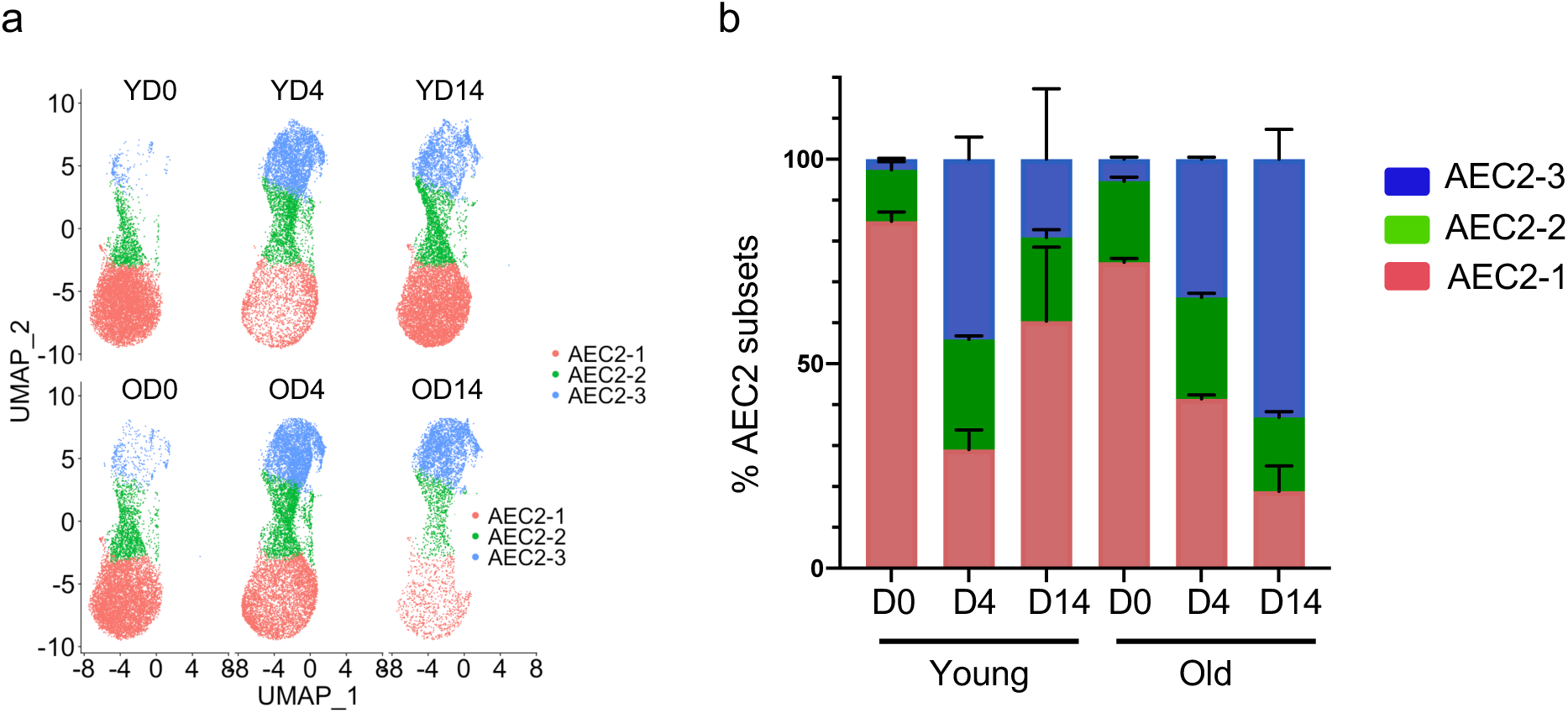
AEC2s clusters were influenced by both aging and lung injury. (a) UMAP showing the distributions of the three subsets of AEC2s grouped by age and injury date (red, cluster1; green, cluster2; and blue, cluster3). (b) Percentage of AEC2 subsets grouped by age and injury date (red, AEC2-1; green, AEC2-2; and blue, AEC2-3).

### IPF AEC2s showed similar gene signatures of murine AEC2-2 and AEC2-3 subsets

We have identified three subsets of AEC2s in mouse model. Subset AEC2-1 represent intact AEC2s and subsets AEC2-2 and AEC2-3 represent damaged AEC2s. Next we tested if our classification of AEC2 subsets was relevant to human disease. We analyzed single cell RNA-seq data of human epithelial cells from lung tissues of IPF patients and healthy donors, and determined the orthologous gene expression in IPF AEC2s. The detail analysis of this single cell RNA-seq data is reported in a separate paper [bioRxiv DOI here]. When compared with AEC2s from healthy donors, IPF lung lost much AEC2s expressing classical AEC2 genes such as surfactant genes including *SFTPC, SLC34A2, ETV5*, and *ABCA3* (Figure 4a). At meantime, we observed strong interferon (IFN) signaling in IPF AEC2s. IPF AEC2s showed higher IFN activation score (Figure 4b) and elevated expression of IFN alpha inducible protein 27 (*IFI27)* (Figure 4c). Growth Arrest Specific 6 (*GAS6*) was significantly upregulated in in IPF AEC2 (Figure 4c). GAS6 mediated macrophage inflammation (Shibata et al., 2014) and Gas6 was increased in IPF fibroblasts (Espindola et al., 2018). The role of IFN genes and GAS6 in AEC2s is unknown.

**Figure 4.**
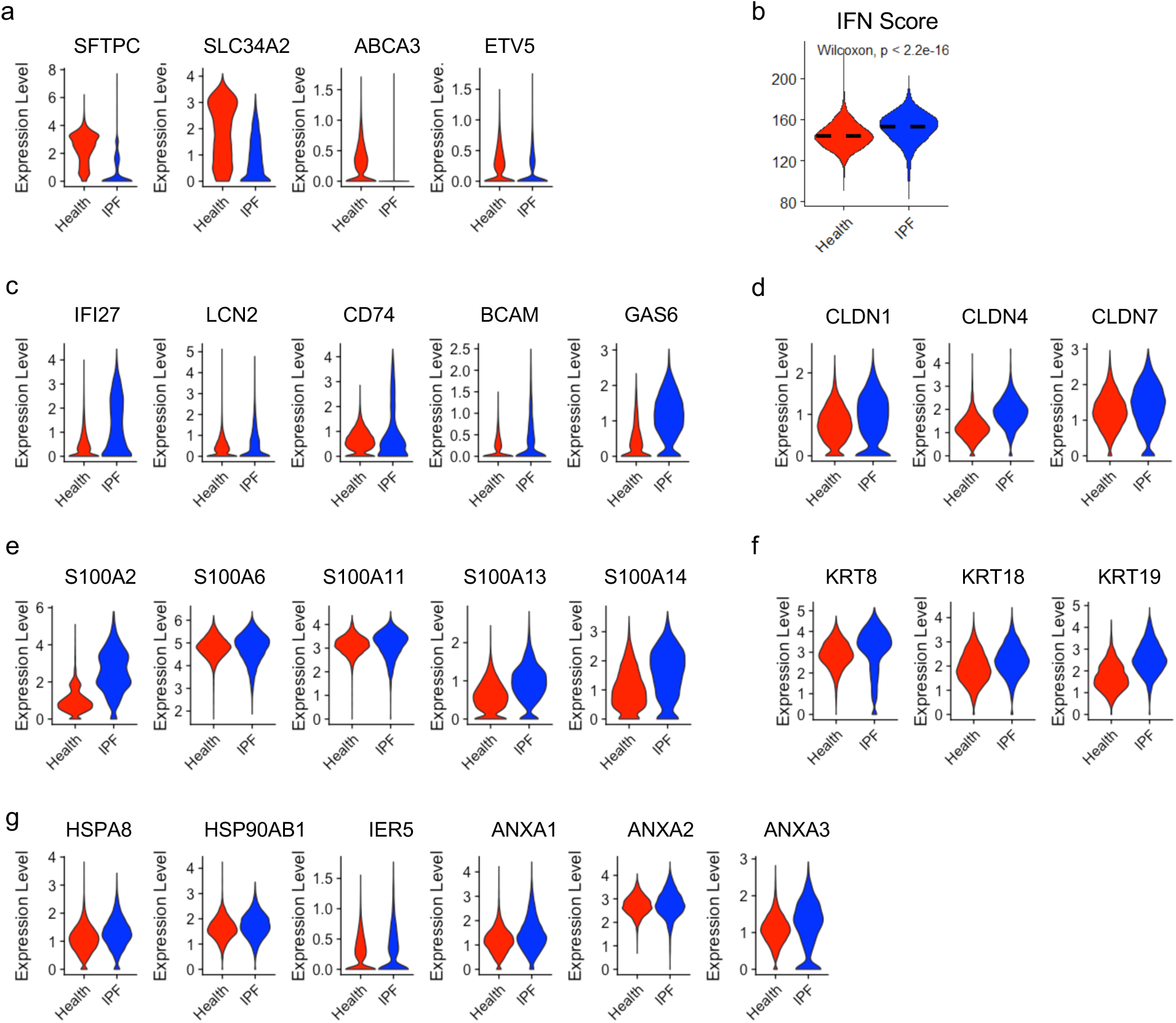
IPF AEC2s share the gene signatures of AEC2-2 and AEC2-3 identified in mouse lungs. (a) Violin plots of expression of mouse AEC2-1 signature genes in IPF and healthy AEC2s (red, healthy; blue, IPF AEC2). (b) Interferon (IFN) activation score of AEC2s from IPF and healthy human lungs (red, healthy; blue, IPF). (c) Violin plots of expression of mouse AEC2-2 signature genes in IPF and healthy AEC2s (red, healthy; blue, IPF AEC2). (d-g) Violin plots of expression of cluster 3 AEC2 signature genes, including CLDN family (d), S100 family (e), Keratin family (f) and Annexin family (g) in IPF and healthy AEC2s (red, healthy; blue, IPF AEC2).

Several claudin family genes (Figure 4d), S100 family genes (Figure 4e), as well as keratin family genes (Figure 4f) were upregulated in IPF AEC2. Both claudins (Kage et al., 2014) and keratins are tight junction proteins and the markers of epithelial differentiation. Recent reports suggest Krt8^+^ or Krt19^+^ AEC2s are progenitor cells in a transitional state from AEC2 to AEC1 in the lung (Kobayashi et al., 2020; Strunz et al., 2020). Much detailed lineage tracing studies and the progenitor characterization especially in IPF are needed to confirm the role of this subpopulation in lung repair.

Genes of heat shock protein family including *HSPA8, HSP90AB1*, and *IER5* were up regulated in IPF AEC2s (Figure 4g). Heat shock proteins have been suggested having a role in pulmonary fibrosis (Bellaye et al., 2018), and they may also contribute to aging by mediating protein quality control (Calderwood et al., 2009). Annexin family genes were upregulated in IPF AEC2 (Figure 4g). These data were consistent with previous reports that IPF AEC2s undergone apoptosis (Korfei et al., 2008).

IPF AEC2s showed the similar gene signature as damaged AEC2s in mouse lung with decreased classical AEC2 marker gene expression, strong interferon signaling, and increased expression of claudin family, S100 protein family, keratin family, heat shock protein family, and annexin family genes. These results suggest an abnormal reprogramming of IPF AEC2s. More detail study is needed to understand how this altered gene expression affects AEC2 function in IPF.

### Lipid metabolic dysfunction of AEC2s in bleomycin-injured old mouse lungs

Lipid metabolism is crucially important for surfactant protein synthesis and maintaining AEC2 functions. Lipid metabolic dysfunction of AEC2s has been linked to aging (Angelidis et al., 2019). We found lower fatty acid biosynthesis (FAB) score of mouse AEC2 subsets AEC2-2 and AEC2-3 relative to AEC2-1 cells (Figure 5a). A group of genes related to lipid biosynthesis and metabolism including *Apoe, Fabp5, Soat1 Scd1, Scd2, Fasn, Cat, Acly, Elovl 1*, and *Ptgs1* were all down regulated in AEC2-2 subset and further reduced in AEC2-3 subset cells (Figure 5b). Furthermore, lipid biosynthesis regulatory genes such *Scref1* (encoding SREBP) and *Insig1* were also down regulated in AEC2-2 and more significant in AEC2-3 (Figure 2b).

**Figure 5.**
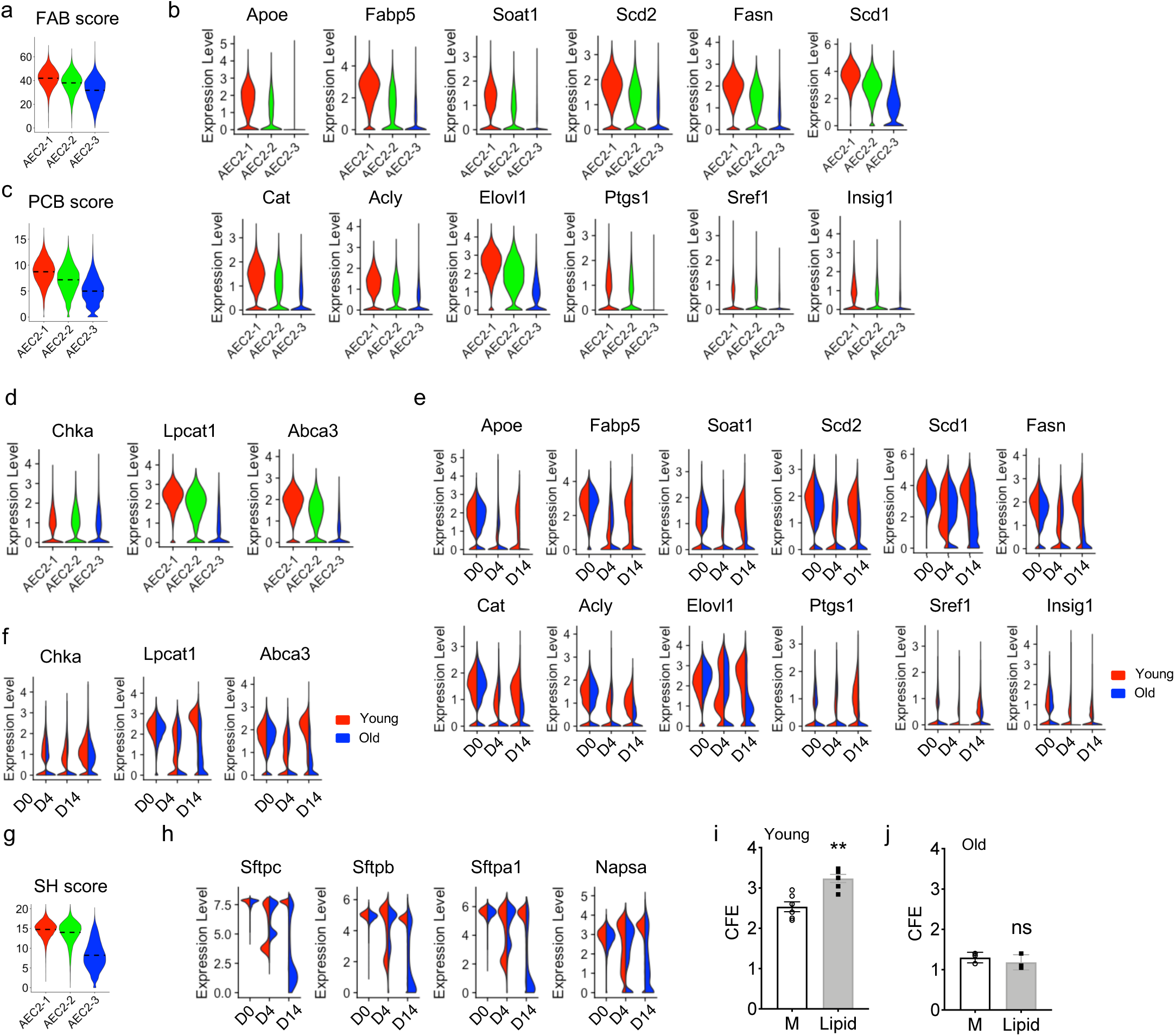
Lipid metabolism of AEC2s in bleomycin-injured old mouse lungs. (a) Activation scores of fatty acid biosynthesis (FAB) of AEC2 subsets. (b) Violin plots of expression of fatty acid biosynthesis-related genes of AEC2 subsets. (c) Activation scores of phosphatidylcholine biosynthesis (PCB) of AEC2 subsets. (d) Violin plots of expression of phosphatidylcholine biosynthesis-related genes of AEC2 subsets. (e) Violin plots of expression of fatty acid biosynthesis biosynthesis-related genes of AEC2s grouped by age and injury date (red, young; and blue, old). (f) Violin plots of expression of phosphatidylcholine biosynthesis-related genes of AEC2s grouped by age and injury date (red, young; and blue, old). (g) Activation scores of surfactant homeostasis (SH) of AEC2 subsets. (h) Violin plots of expression of surfactant homeostasis-related genes of AEC2s grouped by age and injury date (red, young; and blue, old). (i-j). Flow sorted AEC2s (EpCAM^+^CD31^-^CD34-CD45-CD24-Sca-1-) from uninjured young (n = 6, \**\*p* < 0.01) and old mice (j) (n = 3, ns, not significant) were plated for 3D organoid culture in the presence or absence of exogenous lipid mixture. CFE was determined 12 days into the culture. ns, not significant.

Phosphatidylcholine biosynthesis was reduced in AEC2-2 and further reduced in AEC2-3, showing as reduced Phosphatidylcholine biosynthesis (PCB) score of the cells (Figure 5c). A group of genes related to phosphatidylcholine biosynthesis and transport such as *Chka, Lpcat1, Abca3* and were down regulated AEC2-2 and AEC2-3 subset cells (Figure 5d).

Next we examined the expression of lipid biosynthesis and metabolism related genes of murine AEC2s at different timepoint after bleomycin injury. All those genes that we have found highly expressed in subset AEC2-1 and down regulated in subsets AEC2-2 and AEC2-3 in Figure5b were highly expressed in AEC2s of day 0 intact lung and their expression were decreased at 4 dpi with both young and old mice. The expression was partially recoved in AEC2s from young mice at 14 dpi. However, all of these identified genes were further lost with AEC2s from old mouse lungs 14 dpi (Figure 5e).

Phosphatidylcholine converstion enzyme genes *Chka* and *Lpcat1* showed the same pattern as the lipid biosynthesis and metabolism related genes in Figure 5e, they were significantly reduced in AEC2s from old mouse lungs 14 dpi (Figure 5f). Lamellar body marker *ABCA3* (ATP binding cassette subfamily A member 3) is a major transporter for surfactant lipids (Beers and Mulugeta, 2017) and it was significantly down regulated in AEC2s from old mouse lung 14 dpi (Figure 5f).

The dysfunction of lipid biosynthesis and metabolism would directly affect surfactant proteins of AEC2s. We found that subset AEC2-3 cells had the lowest surfactant homeostasis (SH) score among the three subsets of mouse AEC2s (Figure 5g).

Similar to the expression pattern of lipid biosynthesis and metabolism related genes, surfactant genes includeing *Sftpc, Sftpb*, and *Sftpa1* were highly expression in intact AEC2s from uninjured mice. Their expression was reduced in AEC2s from both bleomycin injured young and old mice. It is interesting that at 14 dpi the expression of surfactant genes of the AEC2s from young mice were recovered, whereas the expression levels of surfactant genes of AEC2s from old mice remained low (Figure 5h). *Napsa* (napsin A) which is an aspartic peptidase to process pro-surfactant proteins in AEC2s (Brasch et al., 2003) also failed to recover in AEC2s from old mice 14 dpi (Figure 5h).

To further investigate the biological role of lipid metabolism in AEC2 renewal, we applied exogenous lipid treatment to 3D orgnoid culture of AEC2s isolated from both uninjured young and old mice. Exogenous lipid treatment was able to promote AEC2 renewal from young mouse lungs (Figure 5i). Interestingly, we did not observe an increase in CFE of AEC2s from old mouse lung with exougenous lipid treatment (Figure 5j), indicating that age AEC2s may have intrinsic changes in proventing them to utilize exougenous lipid.

### Dysregulation of lipid metabolism of IPF AEC2s

Next we investigated the lipid metabolism gene expression of IPF AEC2s and compared to that of AEC2s from healthy control lungs through our single cell RNA-seq data with human lung epithelial cells. IPF AEC2s showed lower activation scores of fatty acid biosynthesis (FAB) and phosphatidylcholine biosynthesis (PCB) (Figure 6a). We observed multiple lipid biosynthesis and metabolism related genes including *CHKA, LPCAT1, SCD, SOAT1, FASN, ACLY, CAT*, and *ACOXL* were all down regulated in IPF AEC2s (Figure 6b). SOD2, a responsive molecule to oxidant stress, was down regulated in IPF AEC2s (Figure 6b). IPF AEC2s also showed lower beta oxidation (BO) score (Figure 6c), suggesting that both lipid synthesis and degradation were dysregulated in IPF AEC2s.

**Figure 6.**
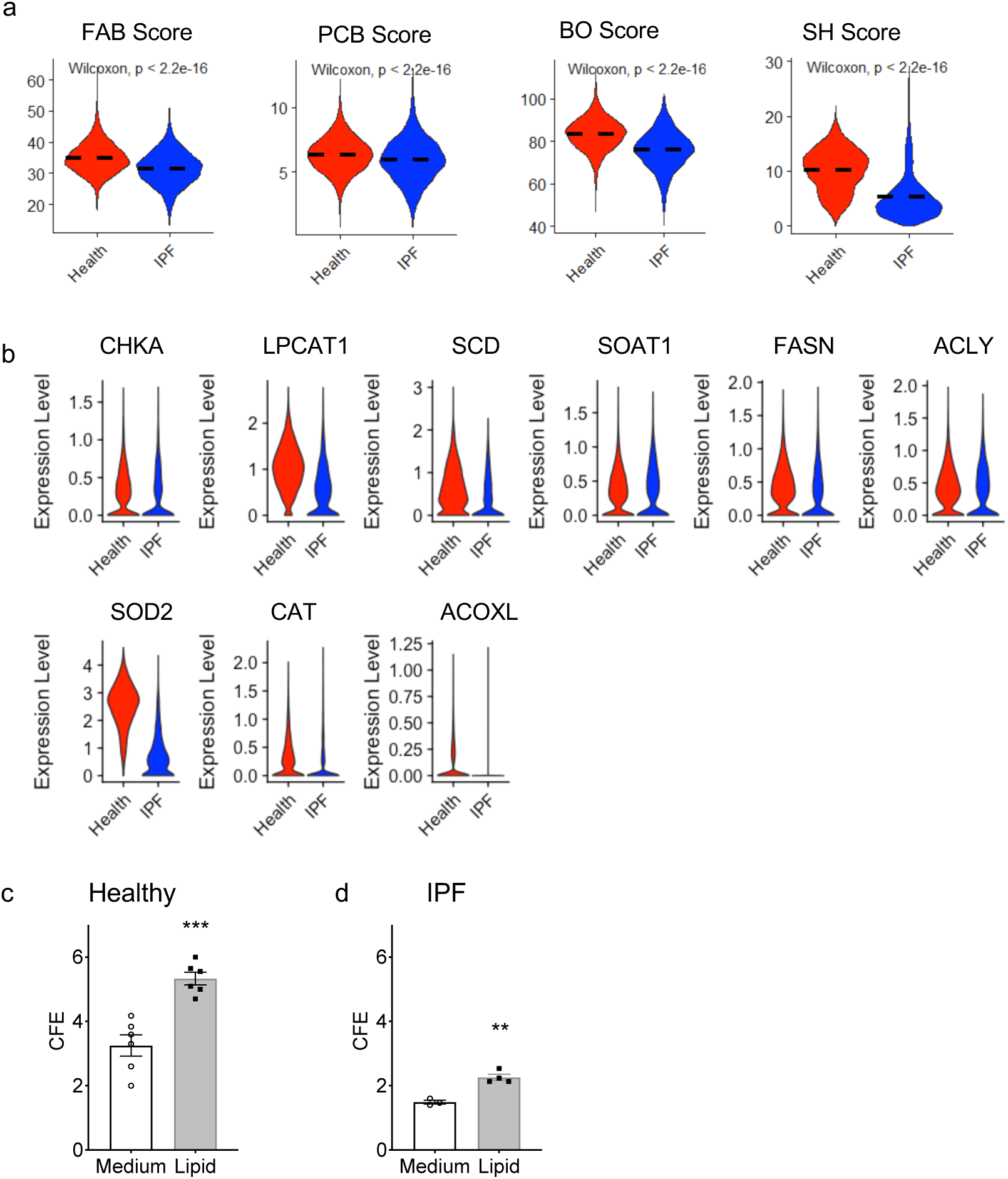
Dysregulated lipid metabolism of IPF AEC2s. (a) Activation scores of fatty acid biosynthesis (FAB), phosphatidylcholine biosynthesis (PCB), beta oxidation (BO), and surfactant homeostasis (SH) of AEC2s from IPF and healthy lungs (red, healthy; blue, IPF). (b) Violin plots of expression of lipid metabolism-related genes of IPF and healthy AEC2s (red, healthy; blue, IPF). (e,f) CFE of flow sorted AEC2s (EpCAM^+^CD31-CD34-CD45-HTII-280^+^) from healthy (e) (n = 6, \*\**\*p* < 0.001) and IPF lungs (f) (n = 3 – 4, \**\*p*< 0.01) in the presence or absence of exogenous lipid mixture.

We have showed that IPF AEC2s have reduced surfactant genes expression relative to AEC2s from healthy lungs (Figure 4a). The down regulation of surfactant genes is assumably resulted from the lipid biosynthesis and metabolism dysfunction of IPF AEC2s. The surfactant homeostasis (SH) score was much lower with IPF AEC2s relative to that of AEC2s from healthy lungs (Figure 6a).

We have reported that there were much fewer AEC2s in the lung of IPF patients compared to that of AEC2s from healthy lung and IPF AEC2s had reduced renewal capacity relative to healthy AEC2s (Liang et al., 2016). We hypothesized that dysregulation of lipid metabolism would contribute to the impaired renewal of IPF AEC2s. As proof-of-principle, we applied lipid treatment to 3D organoid culture of AEC2s from both healthy and IPF lungs. Interestingly, lipid treatment promoted renewal capacity of AEC2s from both healthy and IPF lungs (Figure 6c, 6d).

### Alterated glucose catabolism among AEC2 clusters after injury

Both glucose and lipid metabolism are crutial for AEC2 functions (Beers and Mulugeta, 2017; Lottes et al., 2015). We observed that lipid synthesis genes were reduced in bleomycin injured clusters AEC2-2 and AEC2-3 cells, while genes in the glucose catabolism was enhanced in injured AEC2s (Figure 7a-7c). Both glycolysis score (Figure 7a) and the tricarboxylic acid cycle (TCA) score (Figure 7a) were elevated in subsets AEC2-2 and AEC2-3 cells. Multiple glycolysis related genes includeing *Gapdh, Tpi1, Aldoa, Pgam1, Eno1, Pkm, and Mdh2* were upregulated in cluster AEC2-2, and further elevated in AEC2-3 cells (Figure 7b), suggesting an enhanced glycolysis reprogramming in injured AEC2s.

**Figure 7.**
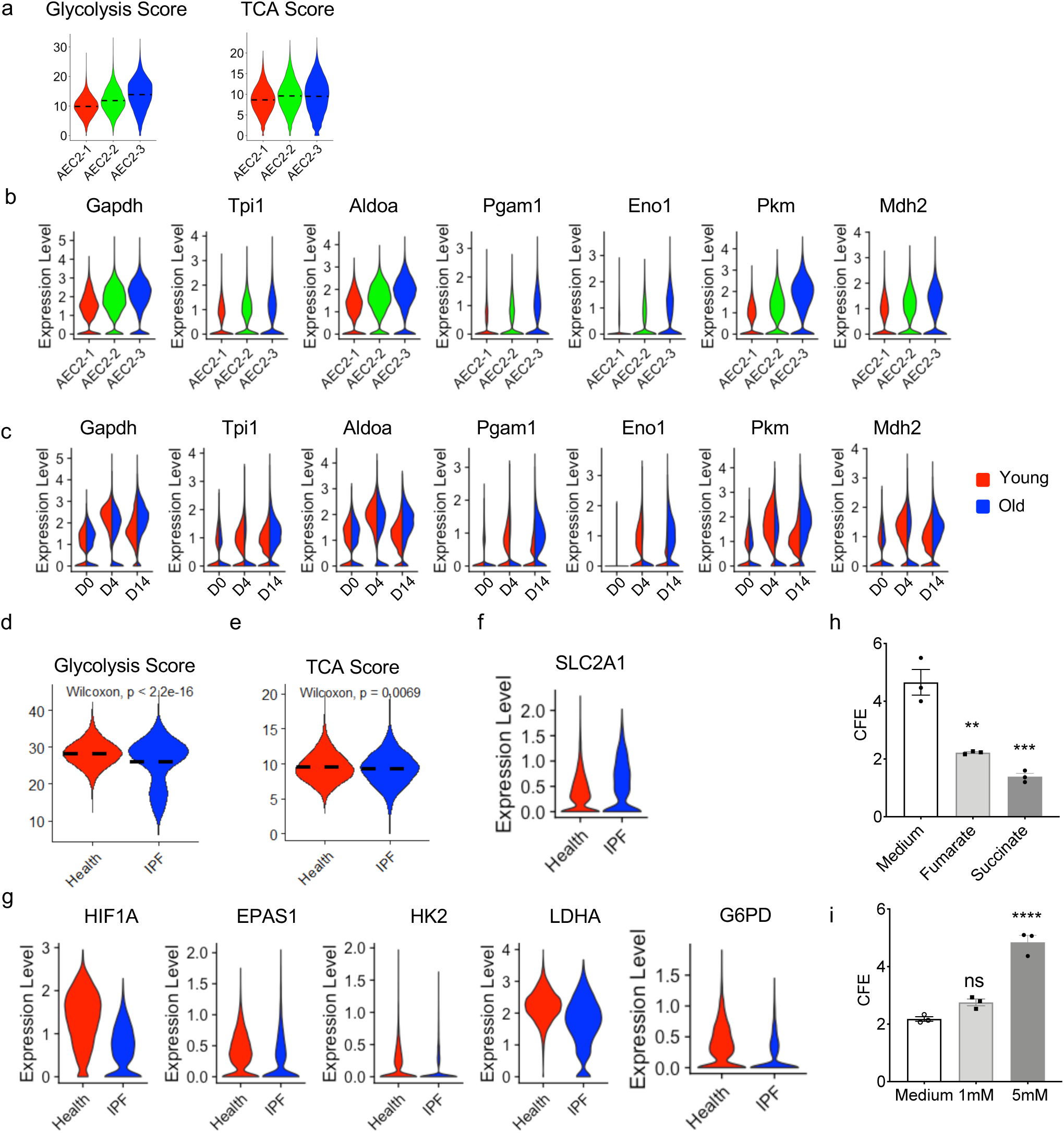
Dysregulated glucose metabolism in aging, injured, and IPF AEC2s. (a) Activation scores of glycolysis and the TCA cycle of the three subsets of AEC2s. (b) Violin plots of expression of glycolysis-related genes of the three subsets of AEC2s. (c) Violin plots of expression of glycolysis-related genes of AEC2s grouped by age and injury date (red, young; blue, old). (d,e) Activation scores of glycolysis (d) and the TCA cycle (e) of IPF and healthy AEC2s. (f) Violin plots of *SLC2A1* expression of AEC2s from healthy and IPF lungs (red, healthy; blue, IPF). (g) Violin plots of expression of HIF1A-related genes of AEC2s healthy and IPF lungs (red, healthy; blue, IPF). (h) TCA cycle intermediates, fumarate and succinate, reduced CFE of AEC2s from uninjured young mouse (n = 3, \**\*p* < 0.01, \*\*\**p* < 0.001). (i) Effect of 2-deoxyglucose (2-DG) at 1 mM and 5 mM on CFE of AEC2s from uninjured young mouse (n = 3, \*\*\*\**p* < 0.0001; ns, not significant).

Furthermore, many of the glycolysis genes were influenced by bleomycin-induced injury and aging. All those genes upregulated in injuried AEC2 subsets shown in Figure 7b were upregulated in AEC2s of bleomycin-injured mouse lungs at 4 and 14 dpi (Figure 7c). It was interesting at 14 dpi, the expression of glucose metabolism related genes of AEC2s from young mice were lowered than that of AEC2s at 4 dpi. However, these gene expression remain high with AEC2s from old mice at 14 dpi.

We did not see significant difference between AEC2s from IPF lung and healthy controls in expression of some of glycolysis-related genes that elevated in AEC2s from injured old mice reported in Figure 7c.

IPF AEC2s from showed slightly lower glycolysis score than that of AEC2s from healthy lung (Figure 7d) and similar TCA score (Figure 7e). We did observe upregulated *SLC2A1* which encodes glucose transport 1 (GLUT1) in IPF AEC2s (Figure 7f). Study showed that Glut1 was upregulated in fibroblasts of aged mice and contributed to lung fibrosis (Cho et al., 2017). To our surprise, *HIF1A, EPAS1* (encoding HIF2A), *HK2, LDHA*, and *G6PD* were all down regulated in IPF AEC2s. It was reported that HIF1A was stablized during acute lung injury (Eckle et al., 2013). Pharmacologic inhibition of HIF1A reduced, while HIF1A activation increased survival of the mice during acute lung injury (Eckle et al., 2013). These data indicate that the glycolysis of AEC2s in IPF lung is dysregulated.

### Impact of glycolysis on AEC2 renewal

To test the biological impacts of glucose catabolism on AEC2 progenitor function, we performed 3D organoid culture of AEC2s isolated from mouse and human lungs with adding substrates or inhibitors of the glucolysis to the culture. Glycolysis substrates, fumarate and succinate reduced colony formation efficiency (CFE) of mouse AEC2s (Figure 7h). On the other hand, inhibition of the glycolysis with 2-deoxyglucose (2-DG) promoted CFE of mouse AEC2s (Figure 7i).

These data suggest that there is a dysregulated metabolic reprogramming including lipid metabolic and glycolysis defects during aging, injury, and fibrosis, and the metabolic reprogramming directly impacts the self-renewal capacity of AEC2s. Therefore, the strategies for restoration of metabolic homeostasis are in favor of AEC2 renewal.

## DISCUSSION

Evidence suggests that IPF is a result of repeated epithelial cell injury and inadequate alveolar epithelial repair (Noble et al., 2012). Aging has been suggested to have a key role in IPF pathogenesis (Raghu et al., 2016; Rojas et al., 2015). Furthermore, metabolic dysregulation has been associated with aging and in patients with IPF (Kang et al., 2016). The molecular links of IPF pathogenesis between aging and metabolism in alveolar epthelial cells are unclear. In this study, we profiled mouse lung epithelial cells after experimental lung injury, identified three subsets of AEC2s during injury and repair, and discovered the metabolic defect in lipid metabolism and glycolysis in aged mouse AEC2s. We further demonstrated that AEC2s from lungs of patients with IPF have the similar gene signature and metabolic dysregulation with injured AEC2s from aged mouse lungs. Perturbation of lipid metabolism and glycolysis significantly affected progenitor renewal capacity of AEC2s.

Heterogeneity of lung epithelial cells has been recognized with recently developed single cell RNA-seq technique (Reyfman et al., 2019; Strunz et al., 2020). Reyfman et al reported subcluters of alveolar epithelial cells from human lung and these clusters distributed differently between fibrotic and healthy donor lungs (Reyfman et al., 2019). In this study we analyzed gene expression of AEC2s from both young and old mice with and without bleomycin injured and identified three subsets of AEC2s. Subset AEC2-1 cells highly express classical AEC2 marker genes and represent intact AEC2s. Majority AEC2s in the uninjured lung and a portion of AEC2s in bleomycin injured mouse lung are AEC2-1 cells. Subset AEC2-2 cells showed strong interferon signaling and highly express Lcn2, Bcam inflammation related genes. Subset AEC2-3 cells highly express keratin and tight junction genes. Subset AEC2-3 cells mainly emerged in the lung after injury. There are very few AEC2s in uninjured lung belong to AEC2-3 subset. Pseudotime analysis suggested a trasition from AEC2-1 to AEC2-2 and further to AEC2-3. A recent study showed a cluster of Krt8 positive AEC2s in bleomycin injured mouse lung are transitional stem cells that precedes the regeneration of AT1 cells (Strunz et al., 2020). The AEC2 in subset AEC2-3 we identified here might include the Krt8+ cells reported. Much detailed lineage tracing studies and the progenitor characterization are needed to confirm the role of this subpopulation in lung repair.

The distribution of the three subsets of AEC2s in the lung is also influenced by aging. At homeostatic stage, old mouse lung contains lower percentage of subset AEC2-1 cells and more AEC2-2 cells relative to that of old mouse lungs, suggesting a chronic AEC2 injury and turnover going on in aged lung without obivious injury. This finding is consistent with reported pro-inflammatory signature of aged lung (Angelidis et al., 2019). At recovery stage after bleomycin injury, ACE2s in the aged lung could not transit back effectively from subset AEC2-2 and AEC2-3 to subset AEC2-1 as it did in the lung of young mice. The failure of transition from damaged AEC2s back to intact AEC2s would be associated with impaired progenitor renewal of AEC2s in aged mouse lung. Increased apoptosis (Korfei et al., 2008), endoplasmic reticulum stress stress (Burman et al., 2018), and senescenece (Minagawa et al., 2011) of aged AEC2s might also contribute to the weakened AEC2 recovery in aged lung after injury. Experimentally, senolytic drugs target alveolar epithelial cells increased epithelial cell markers and attenuate experimental lung fibrosis *ex vivo* (Lehmann et al., 2017).

With analyzing single cell RNA-seq data of human lung epithelial cells, we have found that AEC2s from healthy lung have the similar gene signature with subset AEC2-1 we identified in mice which highly express AEC2 marker genes and is the major AEC2 subset in uninjured lung. On the other hand, IFP AEC2s shared the similar gene signature of subset AEC2-2 and AEC2-3 we identified in injured mouse lungs and further increased in injured old mouse lung. This result indicate that our subset classification of AEC2s with mouse model is bioogicaly relevant to human disease. A recent study reported a cluster of lung alveolar epithelial cells called pre-alveolar type-1 transitional state (PATS) represent cells at transitional stage from AEC2 to AEC1 (Kobayashi et al., 2020). PATS cells highly express CLDN4 and they emerge after injury in mouse lung and are enriched in IPF lung (Kobayashi et al., 2020). In our study, we found that Cldn4 only expressed in subset AEC2-3 cells in injured mouse lung and CLDN4 expression is elevated in AEC2s from IPF lung. The subset AEC2-3 in our study might include the PATS cells reported.

One of the major findings in this study is the identification of metabolic defect in injured and aging AEC2s. We observed that fatty acid biosynthesis, phosphatidylcholine biosynthesis, and the surfactant protein synthesis were all down regulated in AEC2s from bleomycin injured mouse lungs and further reduced with aging. Most importantly we observed down regulation of lipid metabolism related genes in AEC2s from IPF lung. Administration chemically defined lipid mixture to the medium of 3D organoid culture improved colony forming capacity of both healthy and IPF AEC2s. Different effects of lipid metabolism on lung injury and fibrosis have been reported in the literature. Mice with *Elovl1* deficiency died shortly after birth due to epidermal barrier defects (Sassa et al., 2013). Mice with *Apoe* deletion showed impaired alveologenesis, low lung function, and shorter life span compared to wild type mice (Massaro and Massaro, 2008). Lipid synthesis is required to resolve endoplasmic reticulum stress and limit fibrotic responses in the lung (Romero et al., 2018). Targeted deletion FASN (fatty acid synthase) in AEC2s worsened bleomycin-induced lung fibrosis (Chung et al., 2019). We observed significant decrease of FASN expression in IPF AEC2s, consistent with this report. However, Palmitic acid-rich high-fat diet exacerbates experimental pulmonary fibrosis by modulating endoplasmic reticulum stress (Chu et al., 2019). Accumulation of oxidized phospholipids during lung injury (Romero et al., 2015) and aging (Angelidis et al., 2019) contribute to lung fibrosis. Lipids and their regulators have diverse biological functions in lung fibrosis. Some of them prevent and/or repair lung injury while others might induce and/or promote lung fibrosis (Mamazhakypov et al., 2019).

Up regulation of glucolysis has been reported in IPF (Kottmann et al., 2012) and lactic acid induces myofibroblast differentiation(Kottmann et al., 2012; Xie et al., 2015). Serine and glycine synthesis pathway was found necessary for TGF-β-induced collagen synthesis and bleomycin-induced pulmonary fibrosis and PHGDH inhibition attenuated bleomycin induced lung fibrosis (Hamanaka et al., 2018). All these reports indicate glycolysis play a important role in regulating lung fibrosis. In animal model, Elevated Glut1-dependent glycolysis in fibroblasts is associated with the enhanced lung fibrosis of bleomycin injured aged mice and inhibition of glycolysis attenuated lung fibrosis (Cho et al., 2017), Much work has been done in fibroblasts and there is lack of studies with glycolysis reprogramming in AEC2s during aging and in lung fibrosis. We have found increased glucose metabolism related genes in AEC2s subsets AEC2-2 and AEC2-3 which represent inhured AEC2s in mice. AEC2-3 subset showed highest activation score of glycolysis and glucose metabolism related gene expression. Furthermore, AEC2s from bleomycin injured old mouse lung showed persistent high levels of glucose metabolism related genes indicating aging is a important factor regulating glucose metabolism of AEC2s. Unlike gene expression of lipid metabolism which are overlapped between mouse and human AEC2s, the glucose metabolism related genes that we found elevated in injured mouse AEC2s are not upregulated in IPF AEC2s. Instead, IPF AEC2s showed increased expression of GLUT1. Unlike what happened with AEC2s in bleomycin injured aged mice, IPF AEC2s showed lower glycolysis-related genes. The role of HIF1a in lung fibrosis has been reported in fibroblast (Goodwin et al., 2018), macrophages (Philip et al., 2017), as well as in IPF lung tissue (Kusko et al., 2016). We observed that HIF1A expression is decreased in IPF AEC2s. These results suggest that the glucose metabolism of AEC2s in IPF lung is complicated. The mechanism that regulates AEC2 glucose metabolism might not be same between human and mouse. With 3D organoid culture, we demonstrated that glycolysis substrates reduced and inhibition glycolysis promoted mouse AEC2 renewal. Furthere more, inhibition glycolysis promoted renewal capacity of human AEC2s from both healthy and IPF lungs.

In summary, we have identified AEC2 subsets through comprehensive single cell RNA-seq of lung epithelial cells. The gene signatures of AEC2 subsets represent homeostasis and injury stage of AEC2s and they are influenced by aging. Most importantly this AEC2 subset gene signatures are relevant to human AEC2s of IPF. We further indentified dysregulated lipid and glucose metabolism of injured mouse AEC2s and AEC2s from lung of IPF. The aberrant metabolism of injured AEC2s is shown as decreased fatty acid and phospholipid biosynthesis in both injured mouse AEC2s and IPF AEC2s. Increased glycosis was observed with injured mouse AEC2s while the dysregulation of glycolysis in IPF AEC2s is more complicated. Forthermore, aging enhanced the metabolic imbalance in injured AEC2s. Adding lipid mixture to the cells or glycolysis inhibitors promoted AEC2 renewal in 3D organoid culture. Our results indicate restoration metabolic balance of AEC2s with chemical or pharmaceutical reagent might provide therapeutic value for aging related lung injury and fibrosis such as IPF.

## EXPERIMENTAL PROCEDURES

### Animals and Study Approval

All mouse maintenance and procedures were done under the guidance of the Cedars-Sinai Medical Center Institutional Animal Care and Use Committee (IACUC008529) in accordance with institutional and regulatory guidelines. All mice were housed in a pathogen-free facility at Cedars-Sinai. Eight to 12 weeks old (young) and 18 months old (aged) wild-type C57Bl/6J mice were obtained from The Jackson Laboratory and housed in the institution facility at least 2 weeks before experiments.

### Human Lung Tissue and Study Approval

The use of human tissues for research were approved by the Institutional Review Board (IRB) of Cedars-Sinai Medical Center and were under the guidelines outlined by the IRB (Pro00032727), and UCLA Institutional Review Board IRB#13-000462-AM-00019. Informed consent was obtained from each subject.

### Bleomycin instillation

Bleomycin instillation were described previously (Liang et al., 2016). Briefly, animals were randomly allocated to control or treatment groups. Under anesthesia the trachea was surgically exposed. 2.5 U/kg bleomycin (Hospira, Lake Forest, IL) in 25 μl PBS was instilled into the mouse trachea with a 25-G needle inserted between the cartilaginous rings of the trachea. Control animals received saline alone. The tracheostomy site was sutured, and the animals monitored until active. Bleomycin treated mice were actively monitored by trained animal welfare staff until sacrificed. Mice were sacrificed at indicated time points and lung tissues were collected.

### Mouse lung dissociation and flow cytometry

Mouse lung single cell suspensions were isolated as previously described (Liang et al., 2016). In brief, lungs were perfused with 5 ml PBS and then digested with 4 U/ml elastase (Worthington Biochemical Corporation, NJ) and 100 U/ml DNase I (Sigma, St. Louis, MO), and resuspended in Hanks’ balanced saline solution supplemented with 2% fetal bovine serum (FBS), 10 mM HEPES, 0.1 mM EDTA (HBSS+ buffer).

The procedures of staining the cells for flow cytometry and data analysis were described previously (Chen et al., 2012; Liang et al., 2016). In brief, the cell suspension was incubated with primary antibodies including CD24-PE, EpCAM-PE-Cy7, Sca-1-APC, biotinylated-CD31, -CD34, and -CD45, for 45 minutes. Biotin-conjugated antibodies were detected following incubation with streptavidin-APC-Cy7 (catolog # 405208, BioLegend, San Diego, CA). Dead cells were discriminated by 7-amino-actinomycin D (7-AAD) (BD Biosciences, San Diego, CA) staining. Flow cytometry was performed using a Fortesa flow cytometer and FACSAria III sorter (BD Immunocytometry Systems, San Jose, CA) and analyzed using Flow Jo 9.9.6 software (Tree Star, Ashland, OR).

Primary antibodies EpCAM-PE-Cy7 (clone G8.8, Catalog # 118216, RRID AB_1236471) were from BioLegend. CD24-PE (clone M1/69, Catalog # 12-0242-82, RRID AB_467169), Sca-1 (Ly-6A/E)-APC (clone D7, Catalog # 17-5981-82, RRID AB_469487), CD31 (PECAM-1) (clone 390, Catalog # 13-0311-85, RRID AB_466421), CD34 (clone RAM34, Catalog # 13-0341-85, RRID AB_466425), and CD45 (clone 30-F11, Catalog # 13-0451-85, RRID AB_466447) were all from eBioscience (San Diego, CA).

### Human lung dissociation and flow cytometry

Human lung single cell isolation and flow cytometer analysis were performed as described previously (Barkauskas et al., 2013; Liang et al., 2016). In brief, human lung tissues were minced and then digested with 2 mg/ml dispase II, followed by 10 U/ml elastase and 100 U/ml DNase I digestion. Finally, cells were filtered through 100 μm cell strainer, and lysed with red blood cell lysis to get single cell suspension. Antibody staining was similar as with mouse cells. Flow cytometry was performed with Fortesa and FACSAria III flow cytometer and analyzed with Flow Jo 9.9.6 software. Anti-human CD31 (clone WM59, Catalog # 303118, RRID AB_2247932), CD45 (clone WI30, Catalog # 304016, RRID AB_314404), EpCAM (clone 9C4, Catalog # 324212, RRID AB_756086) were from BioLegend. HTII-280 (Gonzalez et al., 2010) was a gift from Dr. L. Dobbs lab at UCSF.

### scRNA-seq and data analysis

scRNA-sequencing was performed in Cedars-Sinai Medical Center Genomics Core. In brief, flow sorted human and mouse single cells were lysed, and mRNA was reverse transcribed and amplified as previously described (Xie et al., 2018). Barcoding and library preparation were performed with standard procedures according to manufacture manuals (10X Genomics, Pleasanton, CA, USA). The barcoded libraries were sequenced with NextSeq500 (Illumina, San Diego, CA, USA) to obtain a sequencing depth of ∼200K reads per cell.

Raw scRNA-seq data was aligned to human genome GRCh38 and mouse genome mm10 with Cell Ranger (10X Genomics), respectively. Downstream quality control, normalization and visualization were performed with Seurat package (Butler et al., 2018). For quality control, the output expression matrix from Cell Ranger was done based on number of genes detected in each cell, number of transcripts detected in each cell and percentage of mitochondrial genes. The expression matrix was then normalized and visualized with UMAP. Silhouette analysis of *k*-means clustering was used to assess the separation distance between the resulting clusters. SCRAT was used to determine and envision high-dimensional metagene sets exhibited in AEC2 subsets as we described previously (Xie et al., 2018). Ingenuity Pathway Analysis (IPA) was performed as described previously (Xie et al., 2018). Differential expression genes with logFC over 0.1 between healthy and IPF AEC2s were analyzed with R software. The data was then imported into IPA software (Qiagen, Hilden, Germany). Canonical pathways from the core analyze were further analyzed. The activation score reflects the sum of expression levels of a biological process (or pathway) related genes in each single cell, the results of each cluster cells were shown in Violin plots. The genes in the biological process (or pathway) were downloaded from UniProt.

### Cell lines

Mouse lung fibroblast cell line MLg2908 (Catalog CCL-206) was from ATCC (Manassas, VA). Mycoplasma contamination was assessed with a MycoFluor™ Mycoplasma Detection Kit (Catalog M7006, Thermo Fisher Scientific) and cells used for experiments were free of mycoplasma contamination.

### 3D Matrigel culture of human and mouse AEC2s

Flow sorted human (Lin^−^EpCAM^+^HTII-280^+^) or mouse (Lin^−^EpCAM^+^CD24^−^Sca-1^−^) AEC2s were cultured in Matrigel/medium (1:1) mixture in the presence of lung fibroblasts MLg2908 cells (Barkauskas et al., 2013; Chen et al., 2012; Liang et al., 2016). 100 μl Matrigel/medium mix containing 3 × 10^3^ AEC2s and 2 × 10^5^ MLg2908 cells were plated into each 24 well 0.4 mm Transwell inserts. 400 μl of medium were added in the lower chambers. Cells were cultured with medium alone or with the treatments indicated. Matrigel (growth factor reduced basement membrane matrix, catalog # 354230) was from Corning Life Sciences (Tewksbury, MA).

For cell treatment, the following chemicals were used. 2% (v/v) chemically defined lipid mixture 1 (catalog # L0288), 100 μM Dimethyl fumarate (catalog # 242926), 5 mM Diethyl succinate (catalog # 112402), 1 and 5 mM 2-Deoxy-D-glucose (2-DG) (catalog # D3179), and 5 μM Phloretin (catalog # P7912) were from Sigma. Same volume of DMSO was used as control.

Fresh medium with proper treatment was changed every other day. Cultures were maintained in humidified 37°C and 5% CO^2^ incubator. Colonies were visualized with a Zeiss Axiovert40 inverted fluorescent microscope (Carl Zeiss AG, Oberkochen, Germany). Number of colonies with a diameter of ≥50 µm from each insert was counted and colony-forming efficiency (CFE) was determined by the number of colonies in each culture as a percentage of input epithelial cells at 12 day after plating.

### Statistics

The statistical difference between groups in the bioinformatics analysis was calculated using the Wilcoxon Signed-rank test. For the scRNA-seq data, the lowest p-value calculated in Seurat was p < 2.2e-10^−16^. For cell treatment data, the statistical difference between groups was calculated using Prism (version 8.4.3) (GraphPad Software, San Diego, CA). Data are expressed as the mean ± SEM. Differences in measured variables between experimental and control group were assessed by using Student’s *t*-tests. One-way ANOVA followed by Bonferroni’s multiple comparison test or two-way ANOVA followed by Tukey’s multiple comparison test was used for multiple comparisons. Results were considered statistically significant at *P* < 0.05.

## Acknowledgments

The authors thank the members of Noble and Jiang laboratory for support and helpful discussion during the course of the study. This work was supported by National Institutes of Health grants R35-HL150829, R01-HL060539, R01-AI052201, R01-HL077291 (PWN), and R01-HL122068 (DJ and PWN), and P01-HL108793 (PWN and DJ). SCR was supported by European Union’s Horizon 2020 research and innovation programme under the Marie Skłodowska-Curie grant agreement 797209. We thank Dr. L. Dobbs of UCSF providing antibodies for the study.

## Conflict of interest

The authors declare that there is no conflict of interest.

## Author contributions

JL, PWN, and DJ conceived the study. JL performed most of the experiments and analyzed the data. GH analyzed single cell RNA transcriptome data, performed flow cytometry analysis, and prepared figures. JL, GH, XL, CY, ND, YW, and DJ analyzed single cell RNA transcriptome data. GH, XL, FT, NL, AB, TX, and SR took part in mouse, cell culture, and biological experiments. PC, CH, BS, and WCP interpreted data and contributed with comments on the manuscript. SSW and JB provided human samples and interpreted data. JL, PWN, and DJ wrote the paper. All authors read and reviewed the manuscript.

## Conflict of Interest

The authors declare that there is no conflict of interest.

## Data availability

The deposition of the raw data files of the single cell RNA-seq are in progress. The accession numbers for the the RNA-seq analyses and the dataset information will be updated accordingly. R code files used for data integration and analysis are available at https://github.com/jiang-fibrosis-lab.

## Data availability

The single cell RNA-seq are deposited

## Conflict of interest

The authors declare that there is no conflict of interest.

## Ethics approval statement

All mouse experimens were under the guidance of the IACUC008529 in accordance with institutional and regulatory guidelines. The use of human tissues for research were under the guidelines outlined by the IRB (Pro00032727).

## Notes

Funding statement: This work was supported by NIH grants R35-HL150829, R01-HL060539, R01-AI052201, R01-HL077291, and R01-HL122068, and P01-HL108793.

### Competing Interest Statement

The authors have declared no competing interest.

